# LPA_3_: Pharmacodynamic differences between lysophosphatidic acid and oleoyl-methoxy glycerophosphothionate. Biased agonism, two sites

**DOI:** 10.1101/2024.09.03.611104

**Authors:** K. Helivier Solís, M. Teresa Romero-Ávila, Ruth Rincón-Heredia, Juan Carlos Martínez-Morales, J. Adolfo García-Sáinz

## Abstract

Lysophosphatidic acid (LPA) and oleoyl-methoxy glycerophosphothionate (OMPT) increased LPA_3_ phosphorylation; OMPT being considerably more potent than LPA. OMPT was also more potent than LPA to activate ERK 1/2. In contrast, to increase intracellular calcium OMPT was less effective than LPA.

LPA-induced LPA_3_-β-arrestin 2 interaction was fast and robust, whereas that induced by OMPT was only detected at 60 min of incubation. LPA- and OMPT-induced receptor internalization was fast but that of OMPT was more marked. LPA-induced internalization was blocked by Pitstop 2, whereas OMPT-induced receptor internalization was partially inhibited by Pitstop 2 and Filipin and entirely by the combination of both. The data again indicate differences in the actions of these agonists.

When LPA-stimulated cells were rechallenged with 1 µM LPA, hardly any response was detected, i.e., a “refractory” state was induced. However, if OMPT was used as the second stimulus, a conspicuous and robust response was observed. These data again suggest the possibility that two binding sites for these agonists might exist in the LPA_3_ receptor, one showing a very high affinity for OMPT and another, likely shared by LPA and OMPT (structural analogs) with lower affinity.

**One sentence summary:** OMPT, oleoyl-methoxy glycerophosphothionate, a biased agonits interacting with an additional binding site in LPA_3_ receptors.

## 1. Introduction

Lysophosphatidic acid (LPA) is a bioactive lipid constituted by a glycerol backbone esterified with a phosphate group at a primary hydroxyl group and with a fatty acid in the sn-1 or sn-2 positions (Supplementary Fig. S1A). In addition to its metabolic roles, LPA exerts a vast series of actions through interaction with various receptors, such as ion channels and nuclear receptors, and with six G protein-coupled receptors (GPCRs), named the lysophosphatidic acid receptor family (*1*). Such GPCRs are divided into two subfamilies according to their sequence homology. The LPA_1-3_ group belongs to the lysophospholipid receptor family, whereas the LPA_4-6_ group is related to the purinergic receptor family (*1*). Structurally, these receptors have seven transmembrane domains with three extracellular loops and three intracellular loops joining them, an extracellular amino-terminus and an intracellular carboxyl-terminus (*1*).

This work is focused on LPA_3_ receptors, which are mainly coupled to G_αi/o_ and G_αq/11_, participating in diverse signaling events such as phospholipase C activation, calcium mobilization, adenylyl cyclase inhibition, and MAPK stimulation (*2, 3*). LPA_3_ receptors are involved in many physiological events, including embryo implantation, expression of antioxidant enzymes, decrease of apoptosis/survival promotion, proliferation, and neuritic ramification, among many others (*2, 3*). A crucial role of LPA receptors, including the LPA_3_ subtype, has been suggested for many cancer types (*4*). Particularly in ovarian cancer, LPA_3_ expression is considered a poor prognosis marker and a therapeutic target (*5*).

Considering the importance of LPA receptors, it is hardly surprising that many groups have attempted to develop selective agonists and antagonists (see, for example,(*6–8*)). Unfortunately, the pharmacological tools with efficacy, potency, and selectivity for these receptors remain very small, including those for the LPA_3_ subtype. 2S-(1-oleoyl-2-O-methyl-glycerophosphothionate) (OMPT) is a derivative of LPA in which a methoxy group substituted the hydroxyl group of the sn-2 position of the glycerol moiety, and the phosphate was changed to a phosphothionate (Supplementary Fig. S1B). It is a potent agonist, initially described as LPA_3_-selective(*6*). Later studies showed that OMPT and related analogs exhibit activity for different LPA receptors, and they only have a weak selectivity for some LPA receptor subtypes (*9–11*). Despite these selectivity problems, OMPT remains a reference for studies on LPA_3_ receptors.

It has been shown that OMPT efficiently increases intracellular calcium in insect Sf9 cells expressing LPA_3_ receptors but poorly in LPA_2_ receptor-expressing cells. In HEK 293 cells transfected with LPA_3_ receptors, OMPT induces ERK phosphorylation and activates GTP[γ-^35^S] binding to cell membranes (*6*). Studies with distinct phosphotionate agonists showed that these compounds could stimulate transforming growth factor-α shedding in cells expressing distinct LPA_1-6_ receptors (*11*).

OMPT has been reported to induce a large variety of actions in vivo and in cellulo; in most cases, the involvement of LPA_3_ receptors was demonstrated using molecular biological approaches. Among these works are studies that show that these receptors participate in erythropoiesis (*12*). Other actions of OMPT include a reduction of the survival rate of cisplatin-treated A549 cells (*13*), an increase in branch formation in hippocampal neurons(*14*), rescue of mitochondrial homeostasis (*15*), an increase in the survival rate of mice with sepsis(*16*), enhanced injury in kidneys subjected to ischemia/ reperfusion (*17*), and induction of cardiac hypertrophy (*18*). It is clear, therefore, that many actions of OMPT have been studied, but to the best of our knowledge, no systematic study is available comparing a variety of cellular events of LPA and OMPT using the same cells, which in our opinion, is a relevant gap of knowledge.

Using HEK 293 cells, conditionally expressing the LPA_3_ receptors, we recently reported their signaling characteristics and defined various responses, described LPA_3_ receptor phosphorylation sites, and explored their roles through mutants with substitution of these sites (*19, 20*). In this work, we comparatively studied a variety of LPA and OMPT actions, and our data clearly show that these agonists exhibit distinct pharmacodynamic characteristics.

## 2. Results

It has been previously shown that LPA induces LPA_3_ phosphorylation (*20*). In this work, we comparatively studied the effect of LPA and OMPT, in parallel experiments, on this parameter. Both agents increase the phosphorylation state of the receptor to a very similar extent (i.e., 2-fold); however, OMPT (EC_50_ value 10 ± 2 nM, n= 10) was considerably (≈ 27-fold) more potent than LPA (EC_50_ 270 ± 70 nM, n= 10; p < 0.001 vs. OMPT) (Fig. 1).

**Fig. 1.**
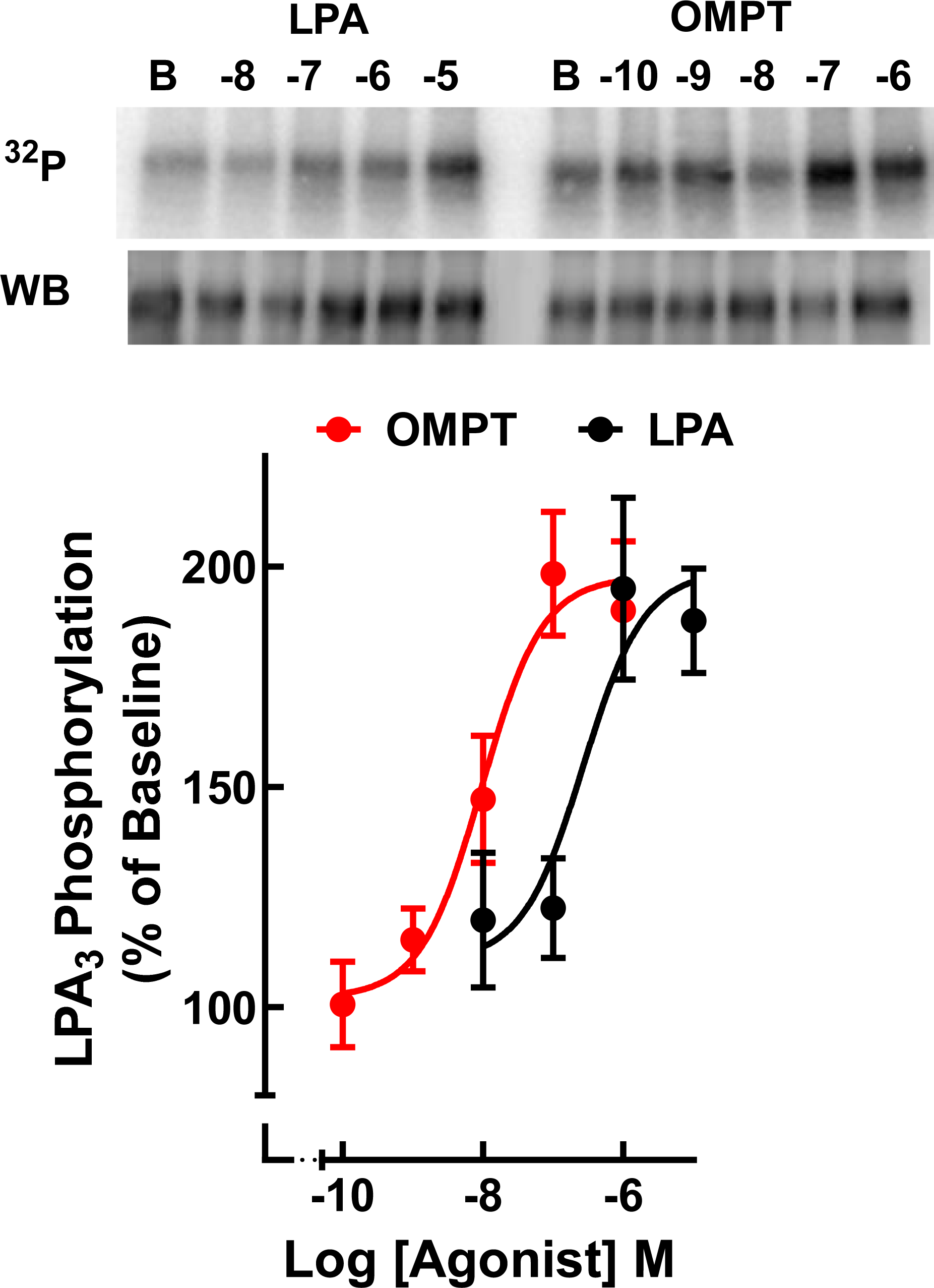
Concentration-response curves for LPA- and OMPT-induced LPA_3_ receptor phosphorylation. Cells were incubated with the indicated concentrations of the agonists for 15 min. Receptor phosphorylation is expressed as the percentage of the baseline value. The means are plotted, and vertical lines indicate the SEM of 10 experiments performed on different days. Representative autoradiographs (^32^P) and Western blots (WB) are presented above the graph.

Similarly, when ERK 1/2 phosphorylation was studied, the efficacies of LPA and OMPT were similar, but again, the potencies were very different; i.e., OMPT (EC_50_ 5 ± 2 nM, n= 6), LPA (EC_50_ 290 ± 50 nm, n=6: P < 0.001 vs. OMPT); i.e., (58-fold difference in potency) (Fig. 2). The time course of LPA and OMPT-stimulated ERK 1/2 phosphorylation showed that LPA and OMPT rapidly increased ERK 1/2 phosphorylation, reaching their maximal at 2 min and progressively decreasing afterward; interestingly, the effect of OMPT was consistently more sustained, and a significant difference was observed at 60 min (Fig. 3). The possibility that different G proteins might be involved in the actions of these agonists was considered, and the effect of pertussis toxin was tested. However, the effect of OMPT was essentially identical in cells pre-incubated overnight without or with pertussis toxin (300 ng/ml) (Supplementary Fig. S2), indicating that the LPA_3_ effect was not mediated through G_i_ but through a pertussis toxin-insensitive G protein, likely G_q/11_, in agreement with data using LPA (*20*).

**Fig. 2.**
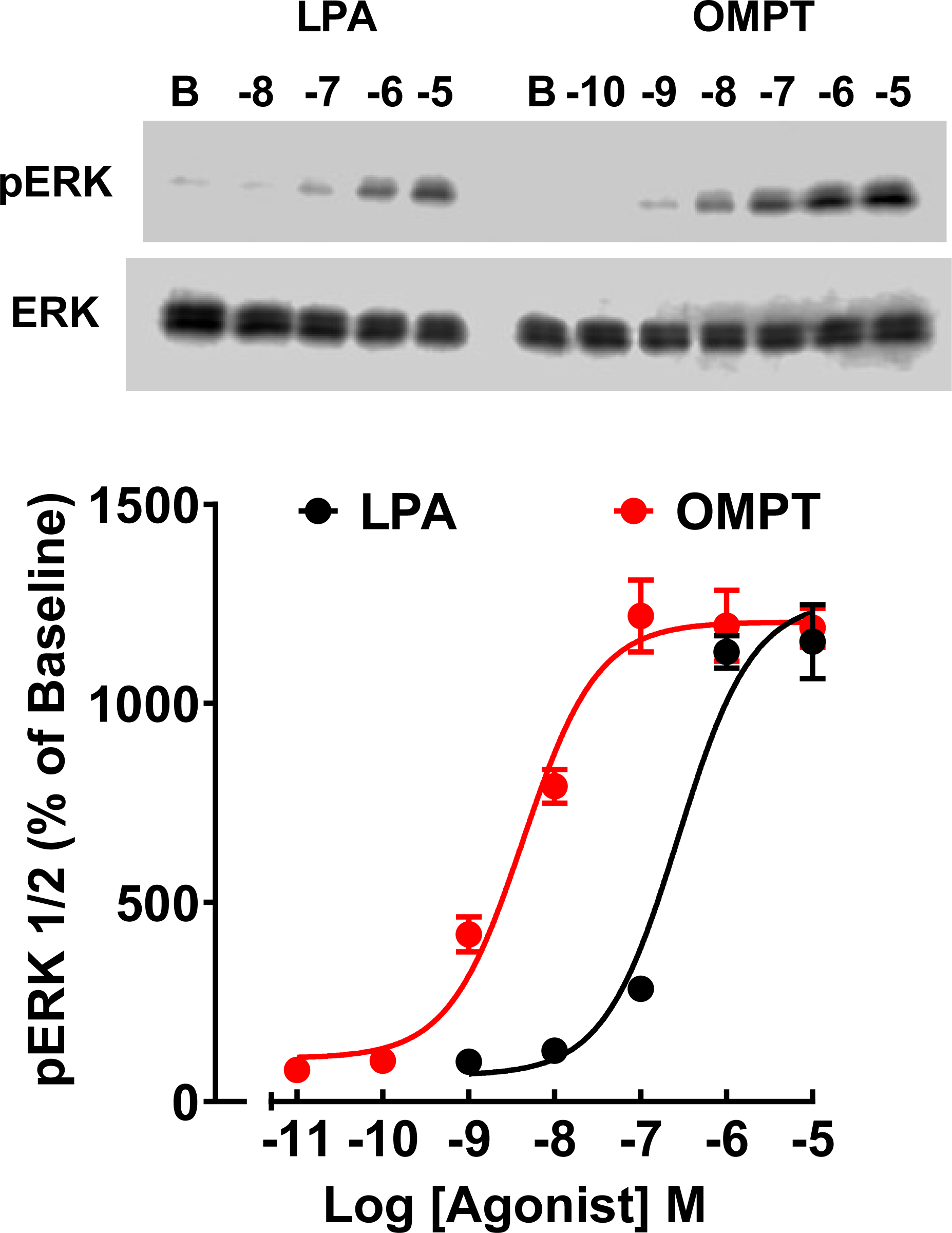
Concentration-response curves for LPA- and OMPT-induced ERK 1/2 phosphorylation. Cells were incubated with the indicated concentrations of the agonists for 2 min. ERK 1/2 phosphorylation is expressed as the percentage of the baseline value. The means are plotted, and vertical lines indicate the SEM of 6 experiments performed on different days. Representative Western blots for phosphorylated (pERK) and total (ERK) kinase are presented above the graph.

**Fig. 3.**
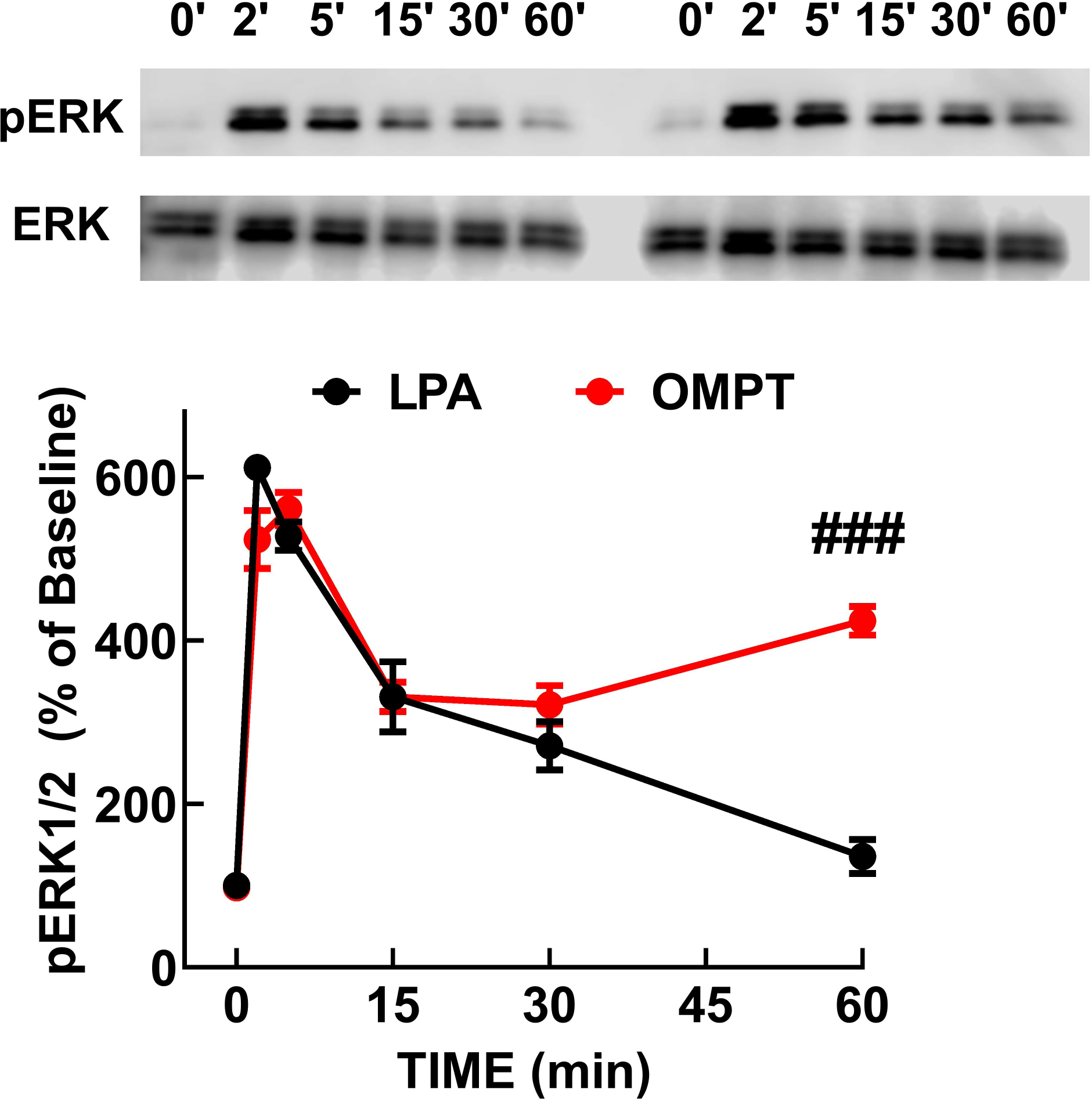
Time course of LPA- and OMPT-induced ERK 1/2 phosphorylation. Cells were incubated for the times indicated with 1 µM of each agonist. ERK 1/2 phosphorylation is expressed as the percentage of the baseline value. The means are plotted, and vertical lines indicate the SEM of 6 experiments performed on different days. Representative Western blots for phosphorylated (pERK) and total (ERK) kinase are presented above the graph. ### p < 0.001 LPA vs. OMPT.

We previously observed that LPA_3_ receptor phosphorylation is associated with its interaction with β-arrestin 2 (*20*). Therefore, we next examined the time course of GFP-tagged LPA_3_ receptor-mCherry β-arrestin 2 biophysical interactions (i.e., FRET, energy transfer, that can only occur if the distance between the proteins is less than 10 nm (*21*)). As expected, when cells were stimulated with 1 µM LPA, a fast (maximum at 2-5 min) and robust increase in FRET was observed that decreased very slowly during the experiment (60 min) (Fig. 4). To our surprise in cells treated with 1 µM OMPT, no significant increase in the FRET signal was observed until the last determination (60 min), when the signal reached levels similar to those observed with LPA (Fig. 4).

**Fig. 4.**
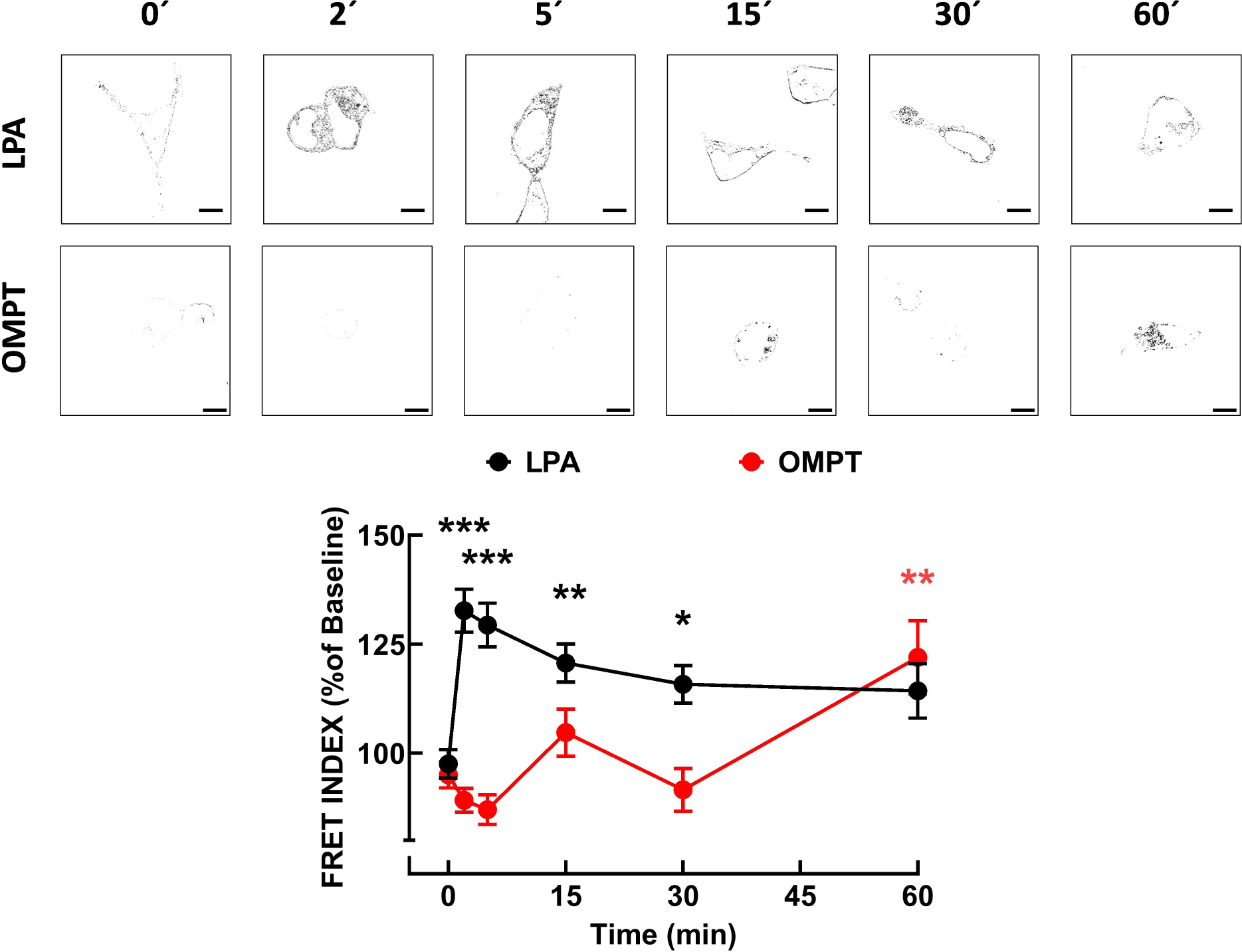
Time-course of LPA- and OMPT-induced LPA_3_-β-arrestin interaction (FRET). Cells were incubated for the times indicated with 1 µM LPA (black symbols and line) or 1 µM OMPT (red symbols and line). The baseline WT FRET index was considered as 100%. The means are plotted, and vertical lines indicate the SEM of 9-10 experiments performed on different days; 10-14 cells were analyzed for each experimental condition in all the experiments. Representative FRET index images are presented above the graph. Bars, 10 µm. *** p< 0.001 vs. baseline, ** p< 0.005 vs. baseline, * p< 0.05 vs. baseline (color coded).

There seems to be a close relationship between LPA_3_-β-arrestin 2 interaction and receptor internalization (*19, 20, 22, 23*). Therefore, LPA- and OMPT-induced LPA_3_ internalization was studied. LPA-induced LPA_3_ receptor internalization was fast, reaching a maximum at 5 min and slowly decreasing at more prolonged incubations. OMPT (Fig. 5A) also induced a fast internalization during the first minutes, but this process increased remarkably later. We have previously shown using this cellular model that although LPA induced internalization, the fluorescent receptors’ delineation of the plasma membrane persisted throughout the incubation. A very discrete, hardly detected at simple sight, but statistically significant decrease in fluorescence abundance was observed during the first 5 min of LPA stimulation (Fig. 5B), which was insignificant afterward. In contrast, it was evidenced that OMPT markedly diminished plasma membrane receptor density, as evidenced by a much lesser fluorescent surface delineation; at more extended times of incubation, plasma membrane fluorescence was discontinuous or poorly defined (Fig. 5B). These differences were striking during continuous observation (see Video legends, Supplementary Video 1 (LPA-induced internalization) and Supplementary Video 2 (OMPT-induced internalization). All these data strongly suggest that different processes could be involved in LPA- and OMPT-induced receptor internalization. In order to test this possibility we took advantage of Pitstop 2, an inhibitor of amphiphysin association of clathrin terminal-domain, which blocks dynamin recruitment (*24*). In Fig. 6A and B, PMA- and OMPT-induced LPA_3_ receptor internalization is shown in cells preincubated for 15 min in the absence (shadowed symbols, dotted lines) and presence of 10 µM Pitstop 2 (solid symbols and lines) before the addition of the agonists. Pitstop 2 markedly inhibited LPA-induced LPA_3_ internalization (Fig. 6A), in agreement with previous findings (*20*) but, in contrast, OMPT-induced internalization was initially delayed by the inhibitor but later increased, reaching values similar to those observed in its absence at 15 and 30 min and without any further increase at 60 min (Fig. 6B) (representative images are presented above the graphs).

**Fig. 5.**
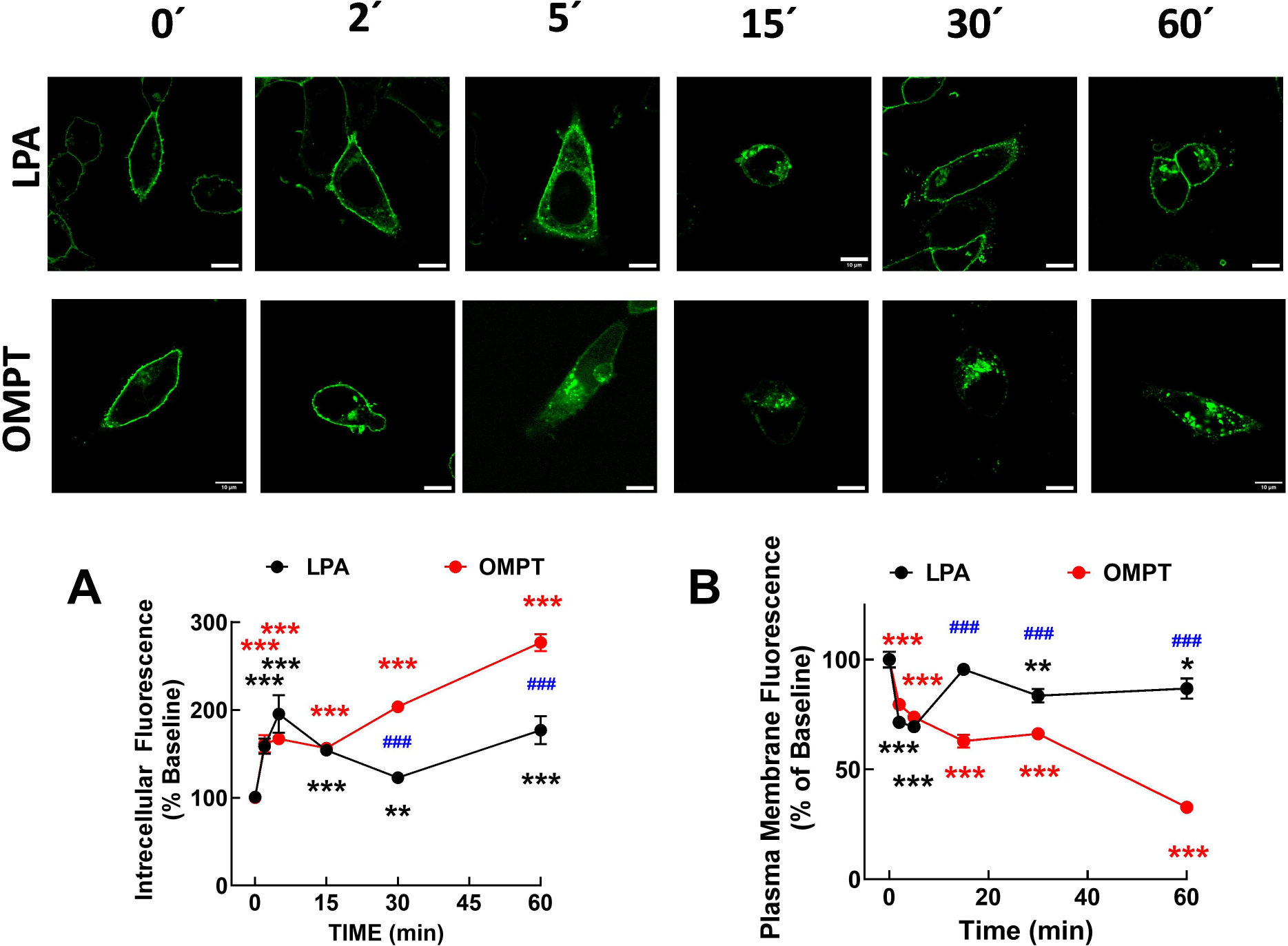
Time course of LPA- and OMPT-induced changes in intracellular (panel A) and plasma membrane (panel B) fluorescence. In both cases, data are presented as the percentage of the baseline values. The means are plotted, and vertical lines indicate the SEM of 4-5 experiments in which 10-14 images were taken for each condition. Representative images (fluorescence, confocal microscopy) are presented above the graph. Bars, 10 µm. *** p< 0.001 vs. baseline, ** p< 0.005 vs. baseline, * p< 0.05 vs. baseline; ^###^ p< 0.001 LPA vs. OMPT.

**Fig. 6.**
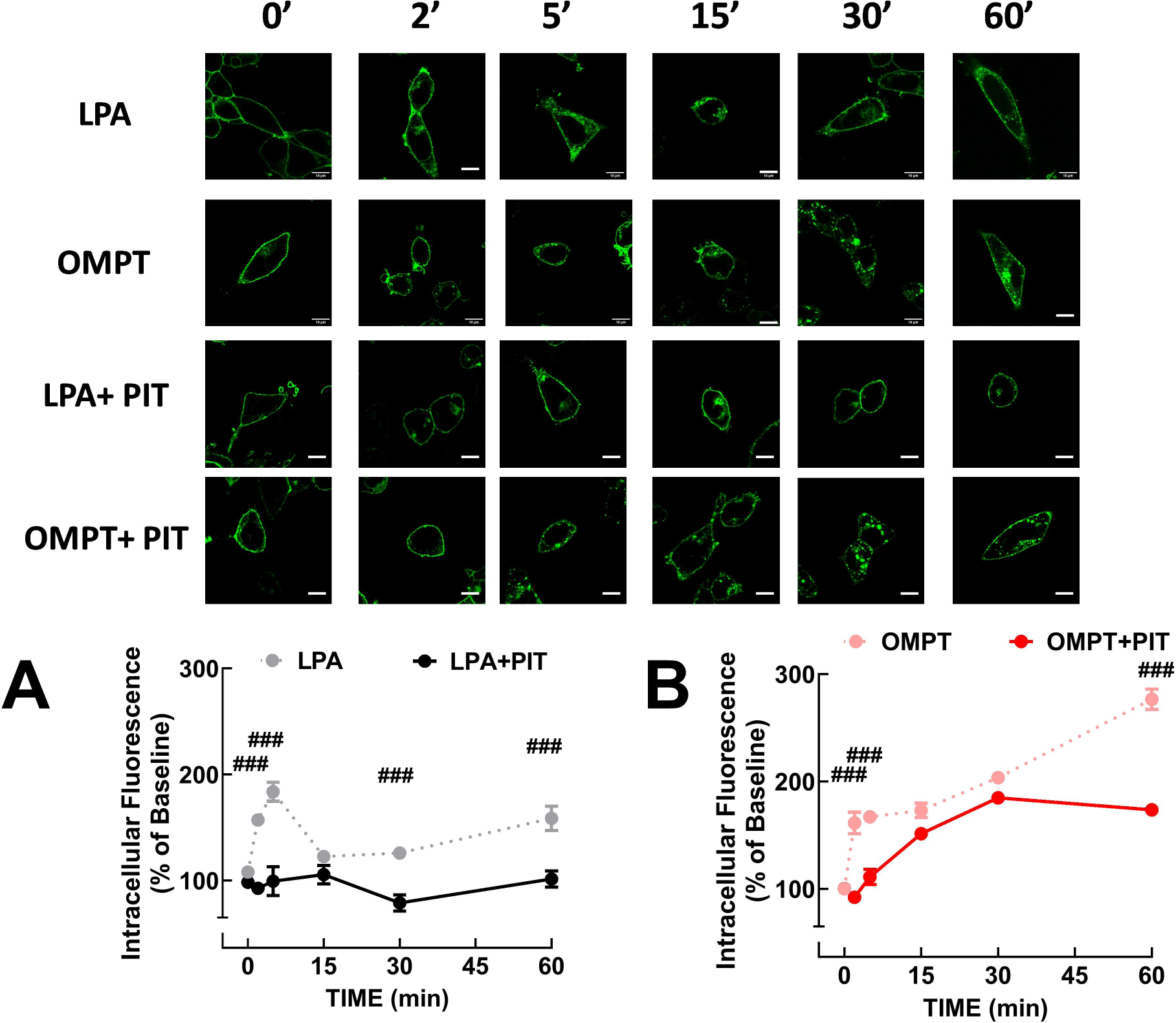
Effect of Pitstop 2 on LPA and OMPT-induced internalization. Cells were preincubated for 15 min without (gray or pale red symbols and lines) or with Pitstop 2 (PIT) (black or bright red symbols and lines) before being stimulated with 1 µM LPA (panel A) or 1 µM OMPT (panel B). The means are plotted, and vertical lines indicate the SEM of 4-5 experiments in which 10-14 images were taken for each condition. Representative images (fluorescence, confocal microscopy) are presented above the graphs. ^###^ p< 0.001 LPA vs. OMPT.

These data are also consistent with the involvement of different internalization mechanisms. To further test this point, we employed Filipin, a sterol-binding polyene macrolide antibiotic (antifungal) inhibitor of the raft/caveolae endocytosis pathway in mammalian cells (*25, 26*). Filipin (1 µM) was preincubated for 60 min. Incubation times were when internalization was evident, i.e., LPA (5 min), OMPT (30 min). PMA (1 µM, 30 min), which also induces internalization, was tested in these experiments because we previously noticed (*19, 20*) similarities with the internalization induced in response to OMPT. Pitstop 2, but not Filipin, reduced baseline internalization (Fig. 7). LPA-induced internalization was essentially blocked by Pitstop 2 (Figs. 6 and 7), as previously reported (*19, 20*), and slightly decreased (not significant) by the Filipin treatment or with the combination of Pitstop 2 and Filipin (Fig 7). In contrast, the action of OMPT was partially inhibited by Pitstop 2 and Filipin and entirely inhibited by the combination of both agents (Fig. 7). PMA-induced internalization was affected by the inhibitors in a way comparable to OMPT, confirming the similarities mentioned above (Fig. 7).

**Fig. 7.**
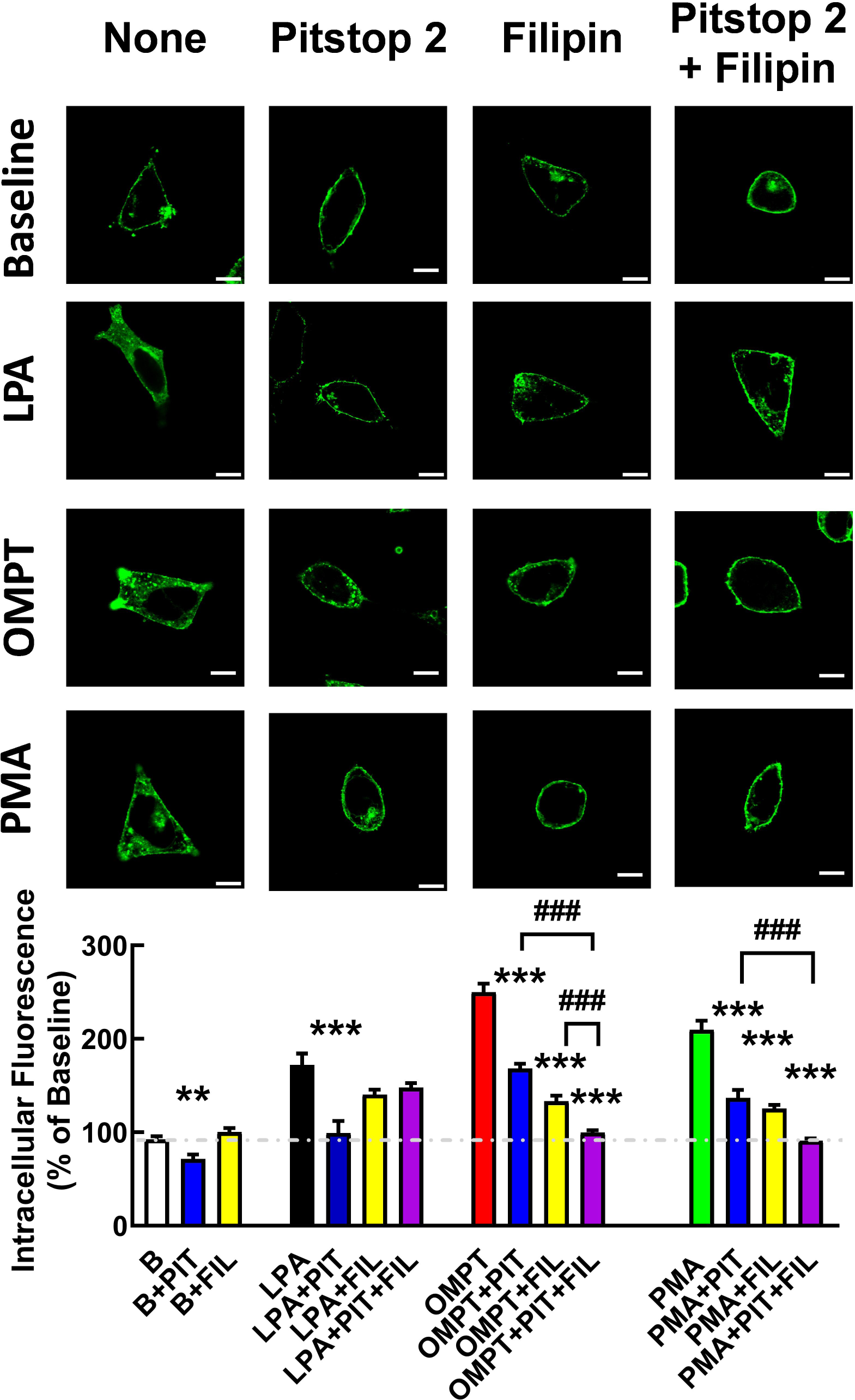
Effects of Pitstop 2 and Filipin on LPA-, OMPT-, and PMA-induced internalization. Cells were preincubated without any internalization inhibitor or with Pitstop 2 (PIT, 15 min, blue columns), Filipin (FIL, 60 min, yellow columns), or both agents (PIT + FIL, purple columns). After the preincubation, the cells were challenged with the agent and for the time indicated: vehicle (B, baseline, 5 min), 1 µM LPA (5 min), 1 µM OMPT (30 min), and 1 µM PMA (30 min). The baseline intracellular fluorescence was considered as 100 %. The means are plotted, and vertical lines indicate the SEM of 5 experiments in which 10-14 images were taken for each condition. Representative images (fluorescence, confocal microscopy) are presented above the graphs.*** p< 0.001 vs. Baseline, ** p< 0.01 vs. baseline; ^###^ p< 0.001, indicated conditions.

Cell proliferation was also tested for a series of agents at concentrations previously observed to be effective (*19, 20*). It is shown in Fig. 8 that serum (10%), LPA (1 µM), PMA (1 µM), OMPT (1 µM), and EGF (100 ng/ml) increased cell proliferation. Interestingly, OMPT was consistently more effective than LPA under these conditions (Fig. 8).

**Fig. 8.**
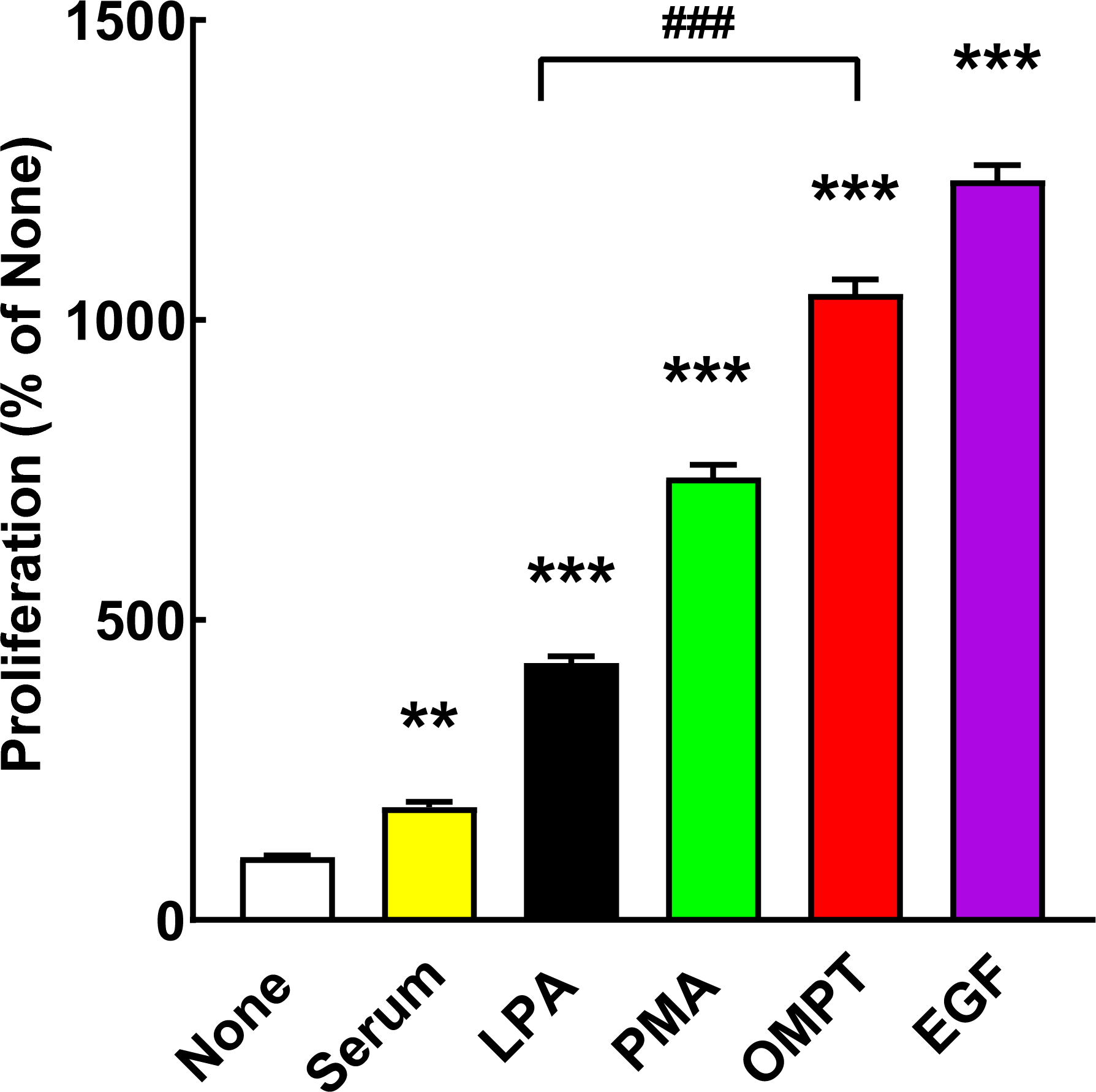
Cell proliferation as reflected by the MTT assay. Proliferation was studied without any agent (None) or with the following stimuli: 10% serum, 1 µM LPA, 1 µM PMA, 1 µM OMPT, or 100 ng/ml EGF. ** p< 0.01 vs. None, *** p< 0.001 vs. None; ^###^ p< 0.001, indicated conditions.

LPA_3_ activation mainly increases intracellular calcium concentration through cation mobilization from intracellular stores (*20*). This parameter was tested by comparing the actions of LPA and OMPT, and the data are shown in Fig. 9. Representative calcium tracings for LPA (panel A) and OMPT (panel B) are presented, and the concentration-response curves are in panel C. It was surprising that OMPT was considerably less effective (OMPT/LPA ratio 0.55), while the potencies of these agents were very similar (LPA 300 ± 40 nM; OMPT 425 ± 50 nM; means ± S.E.M. obtained from 5 curves in each case). The effects of LPA and OMPT on intracellular calcium were also insensitive to pertussis toxin treatment (Supplementary Fig. S3).

**Fig. 9.**
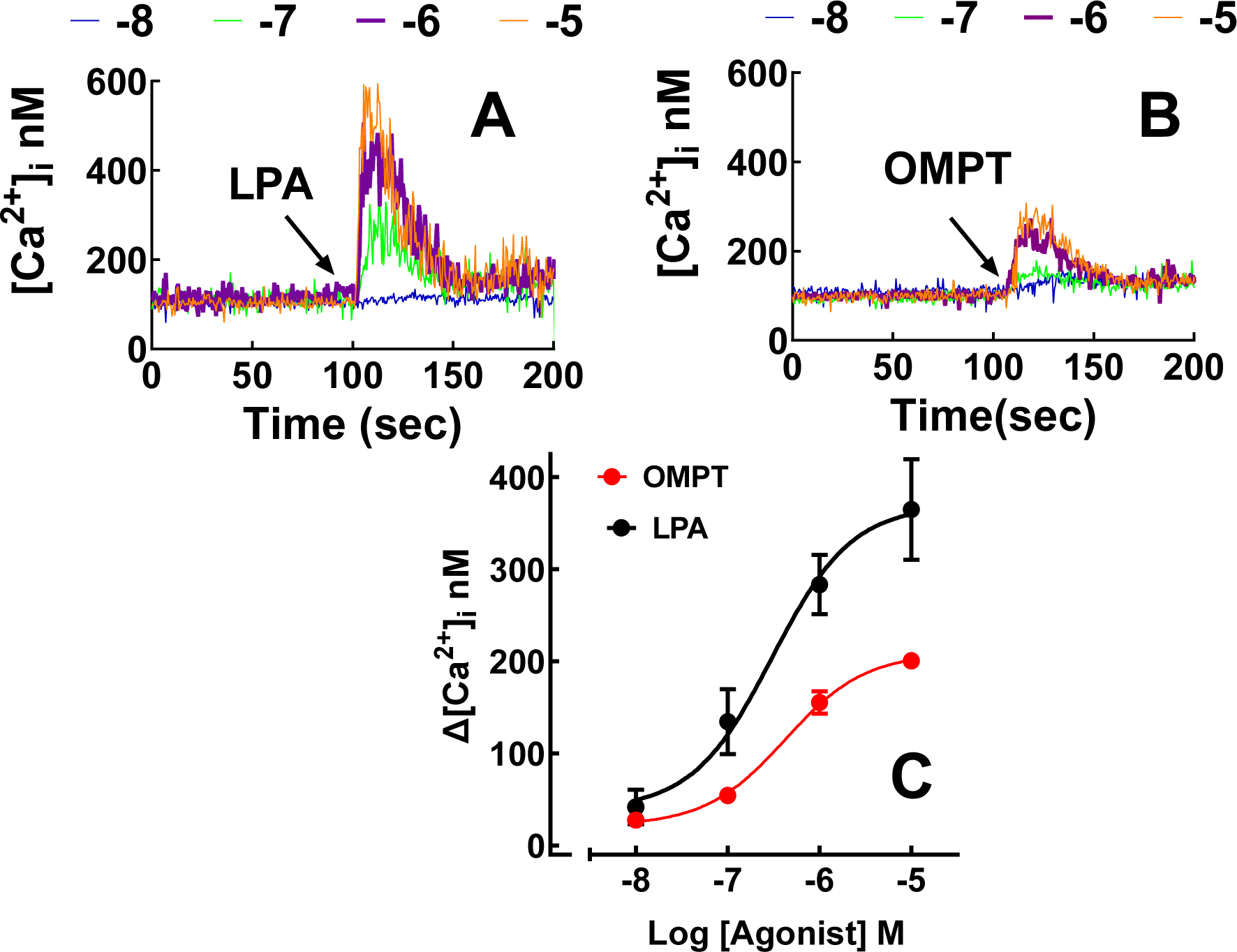
Increases in intracellular calcium in response to LPA and OMPT. Representative calcium tracings of cells incubated with distinct concentrations (color coded) of LPA (panel A) or OMPT (panel B). The concentration-response curves for LPA- and OMPT-induced intracellular calcium increases are presented in panel C. The means are plotted, and vertical lines indicate the SEM of 5-8 distinct curves.

LPA (1 µM) markedly increases intracellular calcium (up to 300 nM, see Figs. 9, 10, and the representative tracing in Fig. 11A). In agreement with previous findings (*19, 20*), we noticed that when LPA-activated LPA_3_-expressing cells were washed and rechallenged with LPA, hardly any homologous desensitization was detected (Fig 10 and a representative tracing in Fig. 11G). Interestingly, when cells were stimulated with LPA, the calcium increase declined, and the cells were rechallenged with 1 µM LPA, hardly any response was detected (Fig. 10; representative tracing in Fig. 11C). This suggested that the continuous presence of LPA induces a receptor’s “refractory state”. Remarkably, when cells were treated with LPA and then challenged with OMPT, a conspicuous and robust response was observed (i.e., surprisingly, no “refractoriness” was detected for this new agonist) as shown in Fig. 10 (representative tracing in Fig. 11D). As shown in Figs. 9 and 10, OMPT (1 µM) also induced a rapid increase in intracellular calcium, but of smaller magnitude (≈ half of PMA action) (representative tracing is shown in Fig. 11B). When cells were treated with 1 µM OMPT, washed and rechallenged again with OMPT, the second response to this agonist was consistently smaller (Fig. 10, representative tracing in Fig. 11J). When OMPT-treated cells were rechallenged (without washing) with this agonist, a diminished response was also detected (Fig 10, representative tracing in Fig 11F), but no evidence of “refractoriness” was detected. These data show that desensitization and “refractoriness” varied with the agonists employed. Cell stimulation with LPA marginally diminished the magnitude of a second stimulation with OMPT when cells were washed between the stimuli (Fig 10, representative tracing in Fig. 11H); a similar effect was observed when cells were incubated with OMPT and rechallenged with the same agent (Fig, 10, representative tracing in Fig. 11J). Incubation with OMPT only slightly diminished the effect of LPA independently of a washing step (Fig. 10, representative tracings in Fig. 11E and Fig. 11I).

**Fig. 10.**
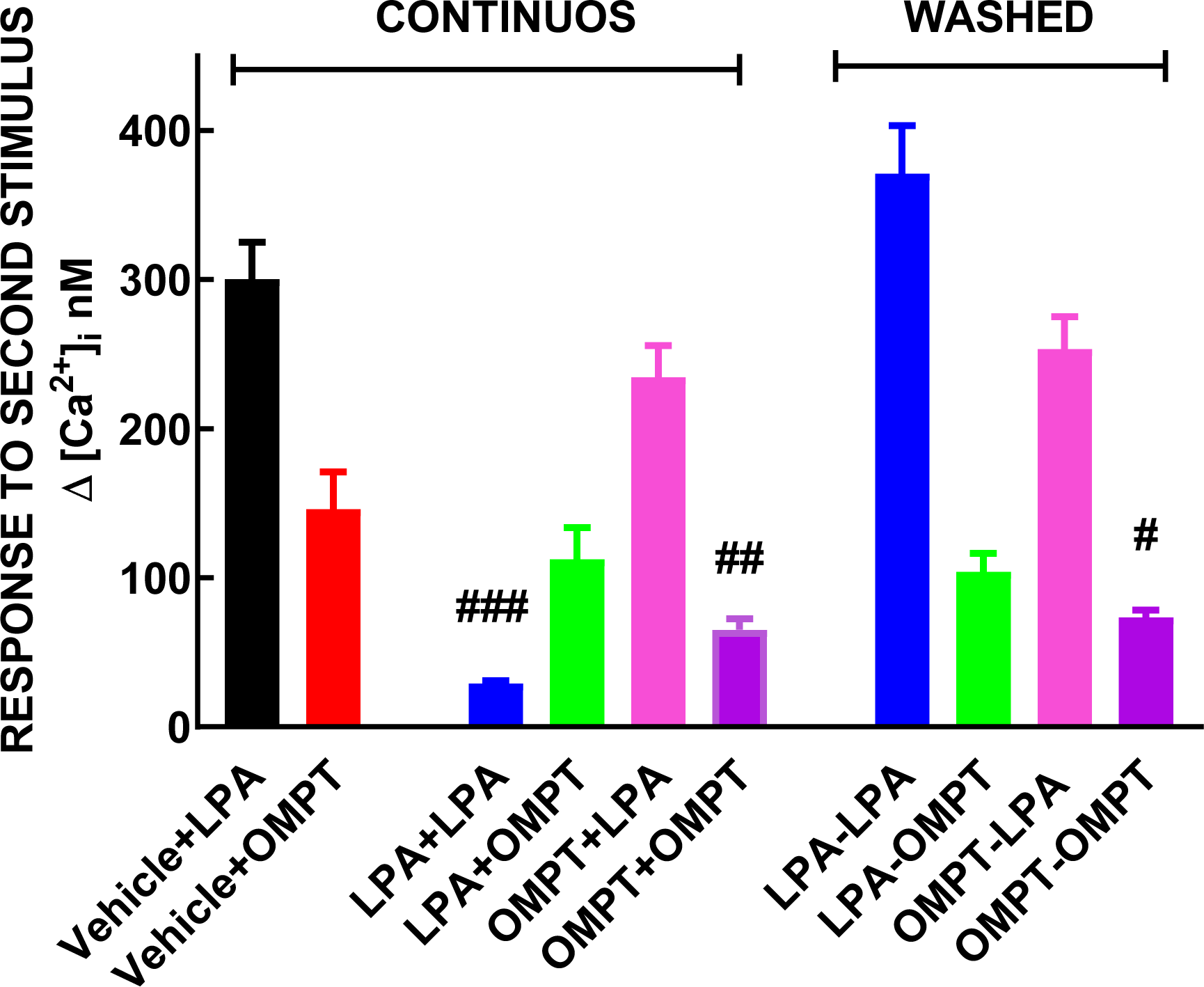
Response to a second stimulation without or with an intermediate washing step. In the first two columns, cells were incubated with the vehicle, followed by a challenge with LPA or OMPT (control responses). In the second group of columns, cells were stimulated with the agonist indicated (first), and when the response vanished, the second stimulus was applied. In the third group of columns, after the cells were stimulated with the first agonist, they were extensively washed to eliminate the agent and rechallenged with the second stimulus. The concentration of LPA and OMPT was 1 µM in all cases. The means are plotted, and vertical lines indicate the SEM of 8-10 determination with cells from distinct cultures. ^###^ p< 0.001 vs. vehicle+LPA, ^##^ p< 0.01 vs. vehicle+OMPT, ^#^ p< 0.001 vs. vehicle+OMPT.

**Fig. 11.**
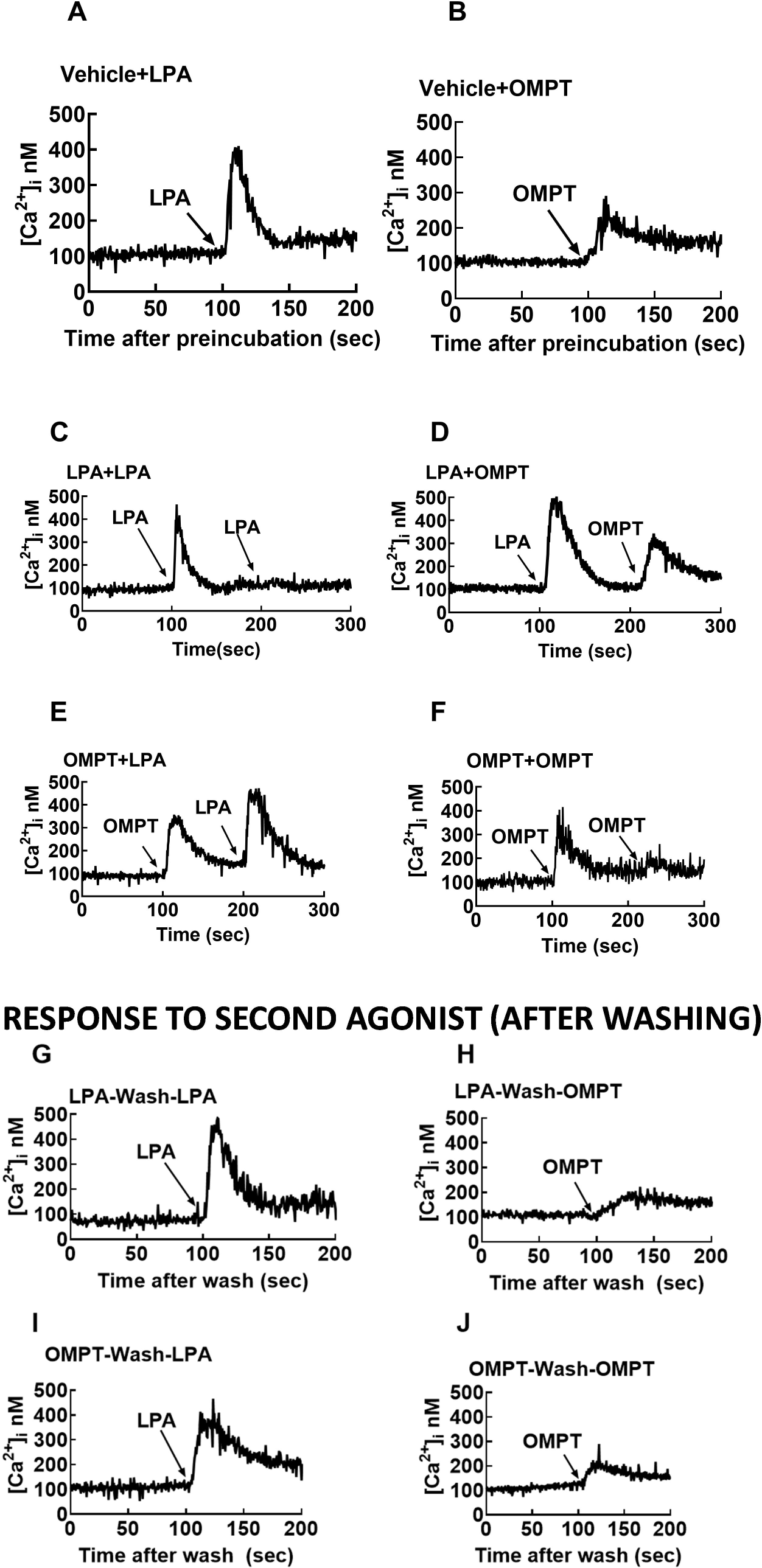
Representative calcium tracings of data that are presented in FIGURE 10.

## 3. Discussion

As indicated in the introduction, other works have studied LPA and OMPT effects in distinct cell types (*12–18, 27–30*), but, to our knowledge, no systematic comparative investigation has been reported. Our present work is a comparative study of various LPA and OMPT actions employing the same cells expressing LPA_3_ receptors, filling a critical knowledge gap.

One of our findings is that OMPT, when compared to LPA, acts as a biased full agonist at low concentrations for some effects (LPA_3_ receptor phosphorylation or ERK 1/2 phosphorylation). In contrast, for other actions, such as the ability to increase intracellular calcium (an almost immediate action closely related to G protein activation), the glycerophosphothionate behaves as a partial agonist, requiring relatively high concentrations to trigger a response. It is also worth mentioning that OMPT-induced ERK 1/2 activation was sustained for longer times than that of LPA; similarly, OMPT-induced LPA_3_ internalization was more intense than that triggered by LPA, particularly at more extended times of incubation (30-60 min) and it was accompanied by a reduction in the plasma membrane receptor density as reflected by cell surface fluorescent delineation. Similarly, 1 µM OMPT induces more intense cell proliferation (as reflected by the MTT test, which indicates cellular oxidative capacity) than an equimolar amount of LPA. Other differences reported in this work include the interaction with β-arrestin 2. It is also interesting that the internalization processes that participate in the actions of LPA and OMPT seem to differ. Our data suggest that LPA-induced internalization occurs mainly via clathrin-coated pits, whereas OMPT action (as well as that triggered by PMA) also involves caveolae.

OMPT exhibits activity in different LPA receptor subtypes, showing only a weak selectivity (*9, 10*). In previous works exploring different parameters, OMPT has been considered a biased LPA agonist. For example, using A549 cells as a model to study LPA_1_ receptors, we observed that OMPT induced a weak increase in intracellular calcium but was able to induce complete ERK 1/2 phosphorylation and cell contraction, suggesting that it behaves as a biased LPA_1_ agonist (*31*). Some antidepressants can activate LPA_1-3_ receptors (*32–35*). It was recently reported that antidepressants and OMPT, acting through LPA_1_ receptors, activate downstream G protein signaling while exerting little effect on β-arrestin recruitment (*10*). Additionally, in the supplementary material of this manuscript, the authors provide evidence that this is also the case for actions mediated through the LPA_3_ receptors (*10*). We show here that OMPT does not mimic the rapid LPA_3_-β-arrestin 2 interaction detected with LPA, but receptor-β-arrestin interaction was only fully detected at later incubation times. The importance of this interaction remains to be studied, but there seems to be some correlation of this process with the intense LPA_3_ internalization and the decreased fluorescent delineation of the plasma membrane observed with OMPT. Although dividing GPCR actions into G protein-mediated and β-arrestin-mediated is appealing, available data suggest that reality is likely much more complex (*36*).

The ability of OMPT to induce receptor phosphorylation is first shown in this manuscript, and it was observed that its potency is much greater than that of the natural ligand, LPA (EC_50_ values 10 nM for OMPT and 260 nm for LPA). Similar, marked differences in potency were also observed for ERK 1/2 phosphorylation (EC_50_ values 5 nM for OMPT and 290 nm for LPA). It was puzzling that OMPT was less effective and showed similar potency to LPA in its ability to increase intracellular calcium (EC_50_ values 425 nM for OMPT and 300 nM for LPA). These data suggested that LPA interacts with a homogeneous LPA_3_ binding site with an apparent affinity of 100-300 nM might represent the orthosteric site. In contrast, OMPT exhibited two potency values, one in the order of 1-10 nm and another with 30-100-fold less potency. The possibility that more than one site might exist for this ligand was considered, i.e., one very high-affinity site not shared with LPA and another likely the common for LPA and OMPT, likely the orthosteric site.

LPA_3_-expressing cells become “refractory” to a second LPA stimulation when it is already present. Such refractoriness was evidenced by the essentially absent response to a second addition of LPA to the media. The possibility that the initial signaling complex (the activated LPA_3_ receptor, G proteins, and possibly other signaling entities) could be trapped in a “frozen” or semi-stable, inactive conformation was considered. It was surprising and puzzling that cells stimulated with LPA and maintained in its presence were readily responsive to OMPT to trigger calcium signaling. OMPT is a partial agonist with a similar apparent affinity to LPA. The data suggest that the stable signaling complex could be disrupted (“relaxed”) by OMPT action; this favors the idea that OMPT might be acting on a different (allosteric) site to induce this; obviously, the possibility that both allosteric and orthosteric sites might participate in the action of this agonist seems likely, but it remains to be experimentally explored. It is worth mentioning that when cells are stimulated and maintained in the presence of OMPT, the addition of LPA triggers a robust response, but if the second stimulus is OMPT, the response is decreased; such decreased second response is not reverted by washing. The data indicate a complex “agonist-selective” response regulation, likely involving different sites and mechanisms. Considering that both LPA and OMPT activate distinct LPA receptor subtypes (*9, 10*), the possibility that a similar two-binding site process might exist for other subtypes is suggested.

We explored the possibility that LPA and OMPT could interact with distinct sites in the LPA_3_ receptor by employing a ligand-receptor docking approach. The data are consistent with this possibility and are presented in the accompanying manuscript.

## 4. Materials and Methods

### 4.1. Reagents

1-Oleyl lysophosphatidic acid (LPA) and 2S-(1-oleoyl-2-O-methyl-glycerophosphothionate) (OMPT) were from Cayman Chemical Co. (Ann Arbor, MI, USA). Phorbol 12-myristate-13-acetate (PMA), as well as Pitstop 2 (N-[5-[(4-Bromophenyl) methylene]-4,5-dihydro-4-oxo-2-thiazolyl]-1-naphthalene-sulfonamide), and Filipin III were obtained from Sigma-Aldrich (St.Louis, MO, USA). Pertussis toxin was purified from vaccine concentrates (*37*). [^32^P]Pi (8500–9120 Ci/mmol) was obtained from American Radiolabeled Chemicals, Inc. (St. Louis, MO, USA). The plasmid for the expression of β-arrestin 2 mCherry-tagged was generously provided by Dr. Adrian J. Butcher (University of Leicester, UK) (*38*). Due to space limitations, the source and catalog number of cells and antibodies and data for other materials are indicated in our previous publications (*19, 20*).

### 4.2 Receptor phosphorylation

It was performed as described (*19*). Briefly, cells were incubated for 1 h in phosphate-free media and 3 h in the same media supplemented with 50 μCi/ml [^32^P]Pi. Labeled cells were treated with the distinct agents, washed, and solubilized in the lysis buffer (*19*). The extracts were centrifuged, and the supernatants were incubated with protein A-agarose and the anti-GFP antiserum generated in our laboratory. Samples were washed, and the pellets were denaturalized with sample buffer (*19*). Proteins were separated using SDS-polyacrylamide gel electrophoresis, electrotransferred onto nitrocellulose membranes, and exposed for 24 h. The amount of phosphorylated receptor was assessed by PhosphorImager analysis using the ImageQuant program. Western blotting for loading controls was performed utilizing a commercial monoclonal anti-GFP antibody.

### 4.3 ERK 1/2 phosphorylation

The cells were serum-starved for 4 h and then stimulated with the indicated concentrations of LPA or OMPT; after this incubation, cells were washed twice and lysed (*19*); the lysates were centrifuged, and proteins contained in supernatants were denatured with Laemmli sample buffer (*39*) and separated by SDS-polyacrylamide gel electrophoresis. Proteins were electrotransferred onto membranes, and immunoblotting was performed. Total- and phospho-ERK 1/2 levels were determined in the same membranes for each experiment; the baseline value was considered 100% for normalization. In all experiments, cell incubations were performed in parallel; the cell extracts were also run together to properly compare the effects of LPA and OMPT.

### 4.4 LPA_3_ receptor-β-arrestin 2 interaction

LPA_3_-GFP construct-expressing cells were transfected with the β-arrestin 2-mCherry plasmid described above (*19*). After 24 h, the cells were collected and seeded on glass-bottomed Petri dishes, and after an additional 48 h in culture, protein-protein interactions were studied. The LPA_3_-β-arrestin interaction was analyzed using Föster Resonance Energy Transfer (FRET), employing an FV10i Olympus microscope. GFP was excited at 488 nm, and emission was detected at 510 nm, whereas mCherry was excited at 580 nm and emitted fluorescence was detected at 610 nm. For FRET channel analysis, GFP (but not mCherry) was excited, and fluorescence was detected at 610 nm; such fluorescence indicated that the proximity among the fluorescent proteins was enough to allow energy transfer (i.e., 1-10 nm) (*21*). The FRET index was quantified using the ImageJ software (version 1.49v) (*40*). The average FRET index obtained with the vehicle (time 0 min) was considered 100%. Individual cells (not clusters) expressing both fluorescent proteins were randomly selected; 10-14 cells were analyzed for each experimental condition.

### 4.5 Receptor internalization

Cells seeded at a low density were cultured on glass-bottomed Petri dishes for 12 h, and LPA_3_ receptor expression was induced with doxycycline (10 µg/ml) (*19*). Before the experiment, the cells were serum-fasted for 1 h. After this incubation, cells were stimulated with LPA or OMPT for the times indicated. A preincubation of 15 min was used to test the action of the clathrin inhibitor (Pitstop 2), whereas a 60 min preincubation was used with the caveolae inhibitor (Filipin); after preincubation, the agents to be tested were added. Cells were washed and fixed as described (*19*). GFP was excited at 488 nm and emitted fluorescence registered at 515-540 nm. The plasma membrane was delineated using the differential interference contrast images to determine receptor internalization. Each cell’s intracellular fluorescence (i.e., excluding the plasma membrane) was quantified as “integrated density”, employing the ImageJ software (*40*), as described(*19*). The baseline intracellular/total fluorescent ratio was considered 100% (the S.E.M. of the different samples in each experiment indicated the internal variation). Plasma membrane fluorescence [1-(intracellular/total fluorescent ratio)] was determined; the baseline value was considered 100 %. Usually, 10-14 images were taken from 3 or 4 cultures obtained on different days for each condition.

### 4.6 Video experiments

The video experiments employed a confocal Zeiss LSM800 microscope with a temperature- and atmosphere (CO_2_ and humidity)-controlled chamber as described (*19*). Excitation, emission, and recording details were as described (*19, 20*). The fact that cells and organelles move (i.e., migrate and change their form) during the experiments and can enter and leave the plane of observation and that LPA and OMPT induce cell contraction and migration should be reminded (*19, 20*).

### 4.7 Cell proliferation

Proliferation was determined using the MTT assay (*41*), as described before (*19*). Cells were seeded in 96-well plates at a density of ≈10,000 cells per well, and receptor expression was induced for 12 h. After induction, cells were treated with the agents indicated for 16 h, and the assay was performed as reported before (*19, 20*).

### 4.8 Intracellular calcium concentration

Determinations were performed as previously described (*19*). In brief, the cells were serum-starved and treated for 12 h with 100 ng/ml doxycycline hyclate to induce LPA_3_ expression. The cells were loaded with 2.5 µM Fura-2 AM for 1 h. Cells were carefully detached from the Petri dishes, washed to eliminate unincorporated dye, and maintained in suspension (*42*). Two excitation wavelengths (340 and 380 nm) and the emission wavelength of 510 nm were employed. Intracellular calcium levels were calculated as described by Grynkiewicz et al. (*43*).

In a series of experiments, cells were subjected to two sequential stimuli to determine if they could respond again. In some of these experiments, the second stimulus was added without removing the first one; in others, after the first stimulus action vanished, the cells were washed in Krebs–Ringer–Hepes containing 0.05% bovine serum albumin (pH 7.4) (*42*) and subjected to the second stimulus.

### 4.9 Statistical analyses

The data are presented as the means ± standard errors of the means. Statistical analysis between two groups was performed using the Student’s t-test and when more groups were compared using ordinary one-way ANOVA with the Bonferroni post-test. Statistical analysis was performed using the software included in the GraphPad Prism program (version 10.2.2). A *P* value < 0.05 was considered statistically significant.

## List of supplementary Material

a. Supplementary Material.doc file
b. Legends for the videos.doc file
c. Video 1 LPA3_LPA.mov and Video 2 LPA3_OMPT.mov files.

## ACKNOWLEDGMENTS

This research was partially supported by Grants from CONAHCYT (Fronteras 6676) and DGAPA (IN201221 and IN201924). The advice and technical support of the following members of the indicated Service Units of our Institute is gratefully acknowledged: Dr. Héctor Malagón and Dr. Claudia Rivera (Bioterio); Juan Barbosa and Gerardo Coello (Cómputo); and Aurey Galván and Manuel Ortínez (Taller). K. Helivier Solís is a student of the Programa de Doctorado en Ciencias Biomédicas (UNAM) of Universidad Nacional Autónoma de México (account 520014983).

## CRediT authorship contribution statement

Conceptualization: KHS, JAG-S

Methodology: KHS, MTR-A, RR-H, JCM-M

Investigation: KHS, MTR-A, RR-H, JCM-M

Visualization: KHS, MTR-A, RR-H, JCM-M

Funding acquisition: JAG-S

Writing – original draft: KHS, JAG-S

Writing – review & editing: KHS, MTR-A, RR-H, JCM-M, JAG-S

## DATA AVAILABILITY STATEMENT

The data are available from the corresponding author upon reasonable request.

## Supplementary Material

**Supplementary Fig S1.**
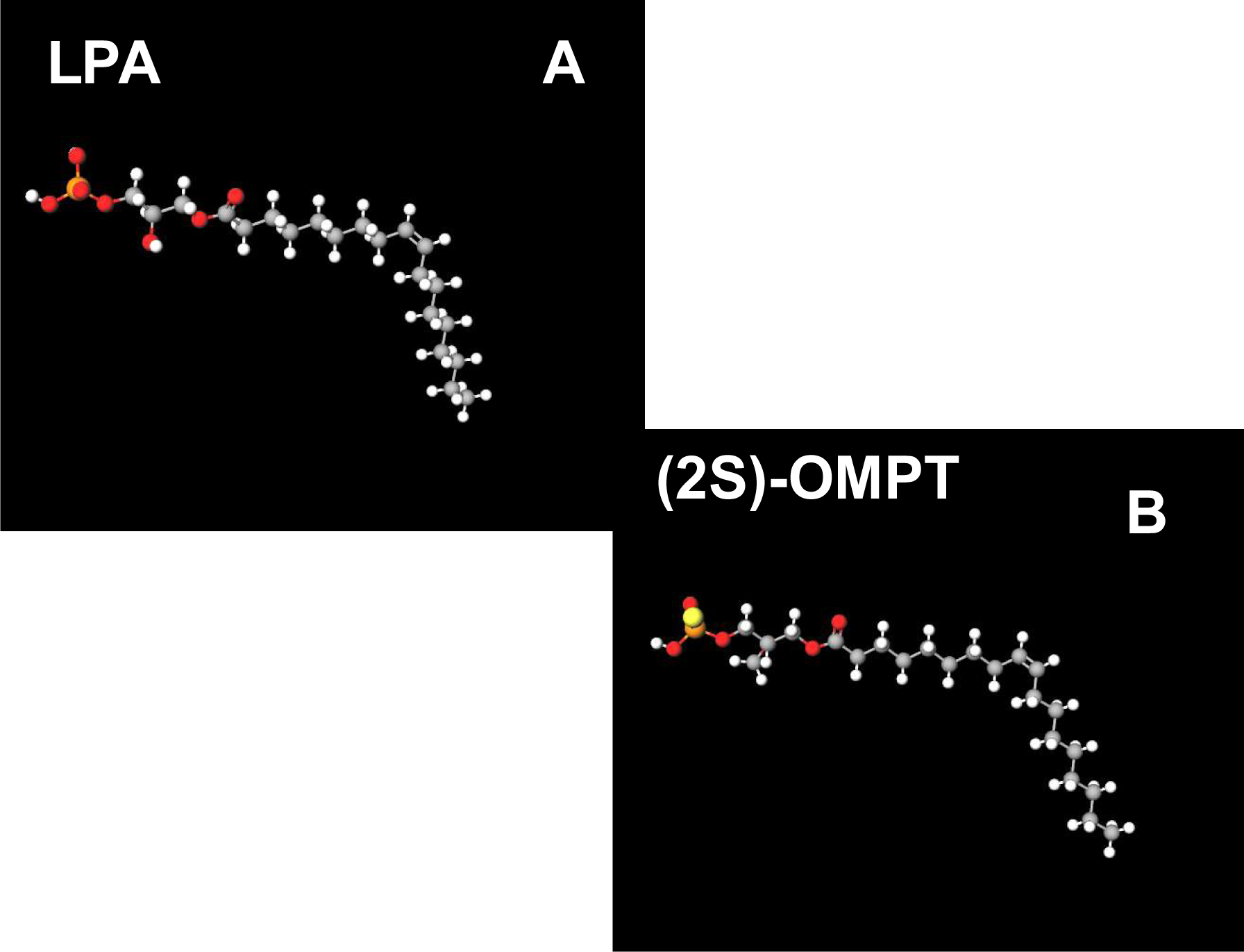
Panel A. Chemical structure of 1-oleoyl-2-hydroxy-sn-glycerol-3-phosphate (LPA). Panel B. Chemical Structure of 2S-(1-oleoyl-2-O-methyl-glycerophosphothionate) ((2S)-OMPT) Atoms in the chemical structure: Carbon (grey), Hydrogen (white), Oxygen (red), Phosphorus (orange) and Sulfur (yellow). https://molview.org/?cid=5497152). Accessed on 04 June 2021.

**Supplementary Fig. S2.**
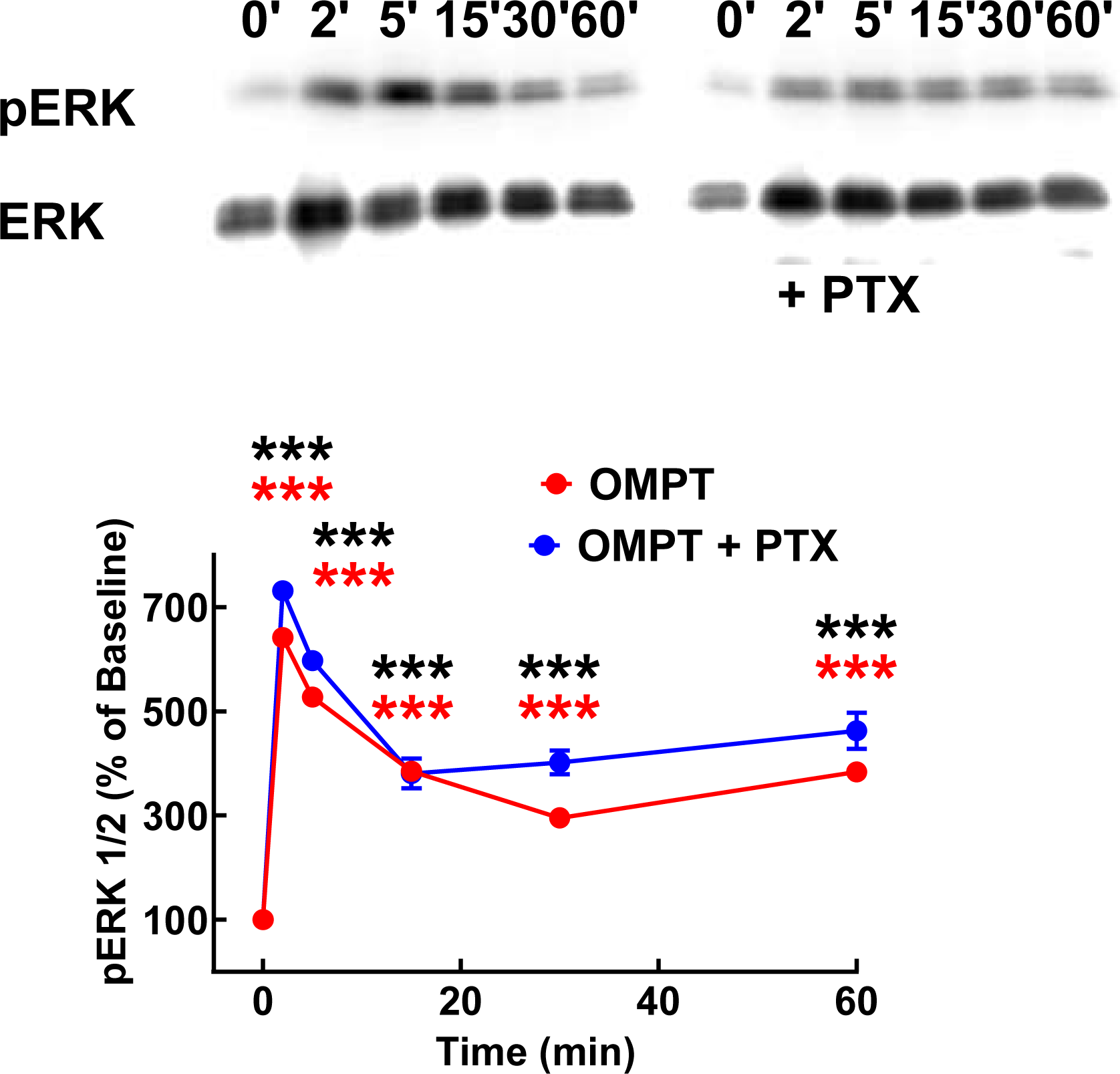
Absence of effect of pertussis toxin on OMPT-induced ERK 1/2 phosphorylation. Cells were incubated overnight without and with pertussis toxin (PTX) and the next day were challenged for the times indicated with 1 µM OMPT. ERK 1/2 phosphorylation is expressed as the percentage of the baseline value. The means are plotted, and vertical lines indicate the SEM of 6 experiments performed on different days. Representative Western blots for phosphorylated (pERK) and total (ERK) ERK are presented above the graph. *** p < 0.001 vs. baseline. Color coded.

**Supplementary Fig. S3.**
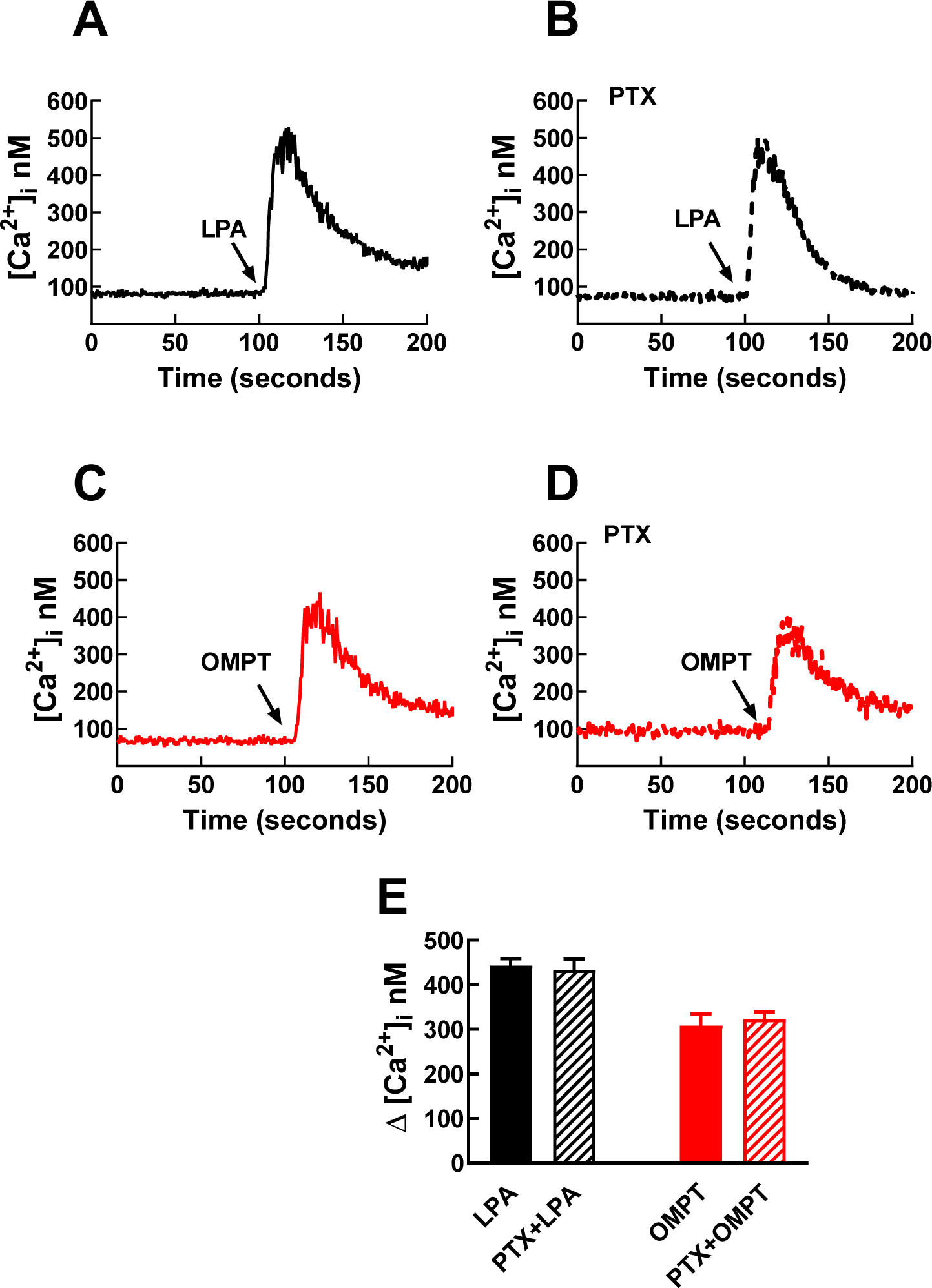
Absence of effect of pertussis toxin on LPA- and OMPT-induced increases in intracellular calcium concentration. Cells were incubated overnight without and with pertussis toxin (PTX); the next day, cells were challenged with 1 µM LPA or 1 µM OMPT. Panels A-D show representative calcium tracings. Panel E shows the increases in intracellular calcium observed. The means are plotted, and vertical lines indicate the SEM of 6 experiments performed on different days.

## LEGENDS FOR THE VIDEOS

**General:** In these studies, recording “Line mode” (1.27 frames per second) was for 5 min (185 frames, no interval) during the baseline, and under-stimulated conditions were for 15 min (556 frames, no interval).

**Video 1**. Effects of LPA. A cell expressing the LPA_3_-GFP construct is shown. Fluorescence is mainly located in the plasma membrane, including filipodia and lamellipodia. Under Baseline conditions, the vesicle traffic is observed. Immediately after adding 1 µM LPA, cell morphology changes (contraction), pearl neckless-like clusters of fluorescence appear, and an increase in rapidly moving vesicles is detected. Only very minor changes in the delineation of plasma membrane fluorescence were detected.

**Video 2.** Effects of OMPT. A cell expressing the LPA_3_-GFP construct is shown. Fluorescence is mainly located in the plasma membrane, including filipodia and lamelipodia but also movement of intracellular vesicles is detected under Baseline conditions. Immediately after adding 1 µM OMPT, the cell contracts and the plasma membrane seems to be markedly decorated with lamellipodia and filipodia (showing a hairy-like fluorescent appearance). Cell contraction continues and blebs randomly appear on the plasma membrane surface. Plasma membrane fluorescence decreases, and its continuity became hard to follow. The cell became full of neckless-like fluorescent structures.

